# Evolutionary constraints in regulatory networks defined by partial order between phenotypes

**DOI:** 10.1101/722520

**Authors:** Manjunatha Kogenaru, Philippe Nghe, Frank J. Poelwijk, Sander J. Tans

**Affiliations:** AMOLF, Science Park 104, 1098 XG Amsterdam, the Netherlands; Laboratoire de Biochimie, UMR CBI 8231, ESPCI Paris, PSL Research University, 10 rue Vauquelin, 75005 Paris, France; Department of Life Sciences, Imperial College London, London SW7 2AZ, United Kingdom; cBio Center at Dana-Farber Cancer Institute, 360 Brookline Avenue, Boston 02215, USA

## Abstract

Gene regulation networks allow organisms to adapt to diverse environmental niches. However, the constraints underlying the evolution of regulatory phenotypes remain ill-defined both theoretically and experimentally. Here, we show that the concept of partial order identifies such constraints, and test the predictions by experimentally evolving an engineered signal-integrating network in multiple environments. We find that populations: 1) expand in fitness space along the Pareto-optimal front predicted by conflicts in regulatory demands, by fine-tuning binding affinities within the network, 2) expand beyond this constraint by changes in the network structure, thus allowing access to new fitness domains. Strikingly, the constraint predictions are based on whether the network output increases or decreases in response to the different signals, and do not require information on the network architecture or underlying genetics. Overall, our findings show that limited knowledge on current regulatory phenotypes can provide predictions on future evolutionary constraints.

## Introduction

Regulatory networks that integrate multiple environmental cues enable organisms to proliferate in diverse environments^1,2^. For instance, bacteria can tolerate highly diverse conditions by recognizing specific combinations of stressors such as pH and osmotic pressure^3^, and plants can elongate above dense canopies by responding to particular combinations of light intensity and wavelength^4^. However, it is not straightforward to establish whether a particular regulatory network is able to optimally respond to the multiplicity of signals presented by a complex environment^5–9^, and consequently how evolutionary constraints limit the range of tolerated environments. Constraints on the adaptation abilities of regulatory networks have been studied both experimentally, by targeted mutagenesis or knock-out of its constituent components^7,10,11^, and computationally, by varying parameters in kinetic models^12–14^. The common denominator for these approaches is their need for detailed information on the network topology and the functioning of its parts, which is lacking for many phenotypes. For instance, evolutionary constraints have been mostly studied for regulatory networks that control developmental programs^15^, but they remain little explored for networks that are involved in competition and selection in variable environments^5,16^. To address these issues, we developed a novel method to identify constraints in signal integration phenotypes that only requires information on current responses, and subsequently tested the predictions using synthetic networks in *E*. *coli*.

## Results

### The order of expressed phenotypes

Central to our approach is *partial order*, which is a concept used in combinatorial optimization problems, such as task scheduling or algorithmic verification^17^, but also in decision making for engineering applications^18^. This concept allows us to categorize signal integration networks that produce monotonic responses to individual input signals — shown to represent a major class of biochemical networks^19^. What we propose is best explained with an example. Consider a phenotype *P*, controlling the fitness of an organism, repressed by an environmental signals *s*_*1*_ and activated by a second environmental signal *s*_*2*_ (Fig. 1a). The vector *S* = (*s*_*1*_, *s*_*2*_) defines the environment and *P*(*s*_*1*_, *s*_*2*_) the response. We consider four environments where these two signals that may be either absent (0) or present (1), and the four corresponding levels of *P* expression (*P*^*00*^, *P*^*10*^, *P*^*01*^, and *P*^*11*^) (Fig. 1b). When switching from an environment *S* = (1,0) to an environment *S* = (0,1), the repressing signal *s*_*1*_ decreases and the activating signal *s*_*2*_ increases. As a consequence of its monotonous response, *P* must increase (from *P*^*10*^ to *P*^*01*^, Fig. 1c). This is the case for any shape of the activation or suppression response, regardless whether its strength or sensitivity is modulated, or how the two signals are integrated, as long as *s*_*1*_ suppresses and *s*_*2*_ activates *P*. Thus, for evolutionary processes that modify any of the activation characteristics of the two signals, one can identify the following constraint on *P*, namely: *P*^*01*^>*P*^*10*^. This indicates that when considering phenotypic responses *P*^*10*^ and *P*^*01*^, what is constrained is their *order*.

**Figure 1.**
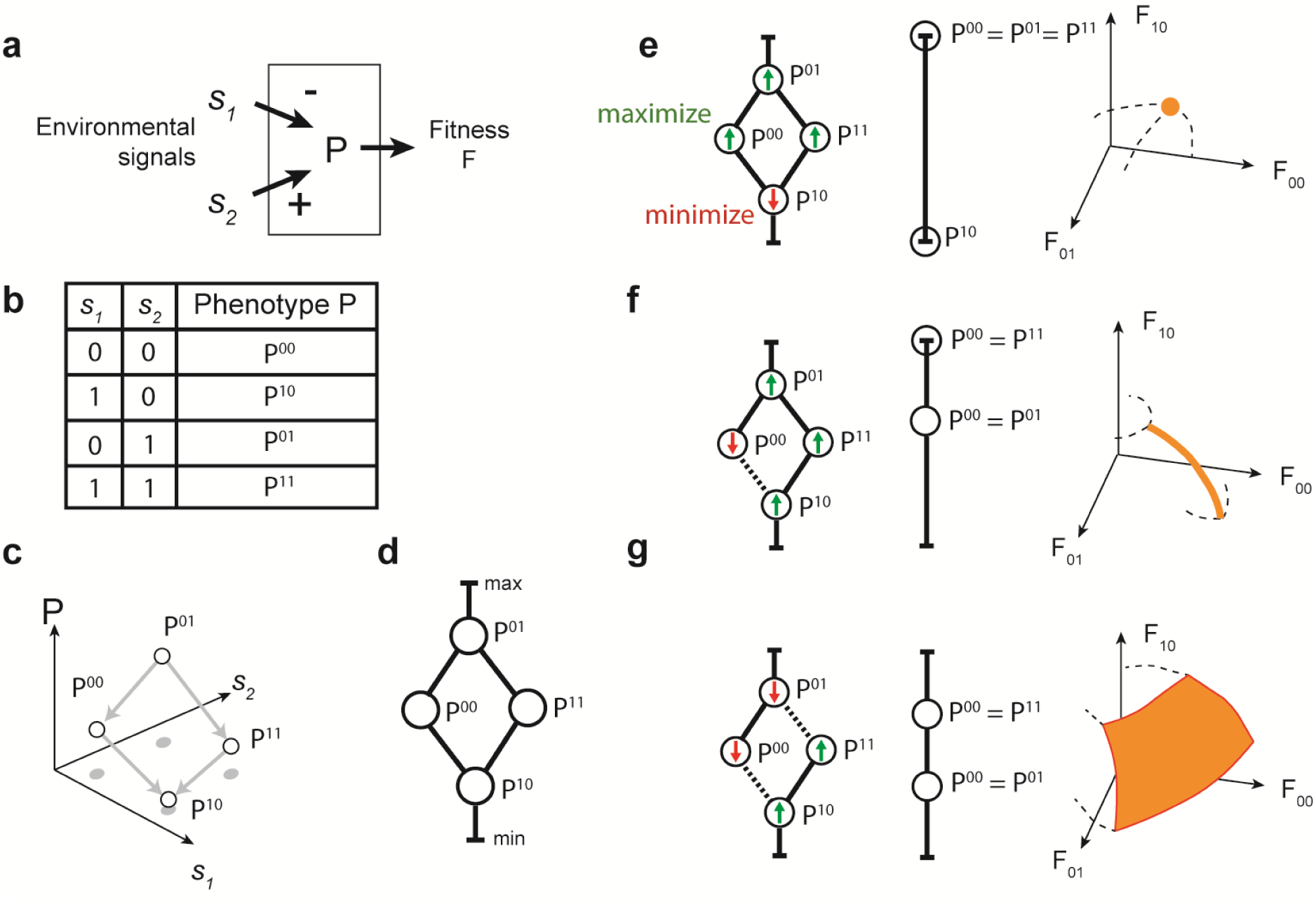
Partial order of phenotypes identify conflicting regulatory objectives and constraint. **a)** Schematic diagram of the system studied. An organism exhibits a phenotype *P* that controls fitness *F*, in response to two environmental signals *s*_*1*_ and *s*_*2*_. Here, expression of *P* is suppressed by *s*_*1*_ and activated by *s*_*2*_. **b)** Considered environmental conditions, which alternate in time. The two signals *s*_*1*_ and *s*_*2*_ can be either present at a particular concentration (1) or absent (0), resulting in four possible values for phenotype *P*. **c)** Diagram indicating the order of expressed phenotypes corresponding to panels a-b. *P*^*01*^ is highest because *P* is not suppressed (by *s*_*1*_) and only stimulated (by *s*_*2*_), while *P*^*10*^ is lowest because *P* is only suppressed and not stimulated. *P*^*00*^ and *P*^*11*^ are both higher than *P*^*10*^ and lower than *P*^*01*^, as they are neither fully stimulated, nor fully repressed. The arrows indicate this order in expressed phenotypes. In contrast, the order between *P*^*00*^ and *P*^*11*^ is not defined because it remains undetermined whether *P* is more activated or more suppressed. **d)** Hasse diagram representing the partial order described in panel c. Phenotypes are represented as nodes, their order by connecting lines. The order is partial because some phenotypes are ordered while others are not. The phenotype *P* can vary between the minimum (min) and the maximum value (max). This partial order is an evolutionary constraint: it remains identical for mutations that change any property of the activation or suppression, as long as the same signals continue to activate or repress. **e-f)** Prediction of conflicts, trade-offs, Pareto-optimal front, and its dimensionality. Consider regulatory evolution driven by mutations and selective pressure. Generally, each of the four alternating environments may select either to increase (maximize) or decrease (minimize) expression of *P*. A partial order analysis predicts constraints in this adaptation. **e)** Set of four selective environments that do not pose a constraint for this response and partial order (depicted in panel c). Given the selection (arrows, left), all nodes can be optimized without conflict (middle), resulting in a single optimum phenotype in fitness space (right, schematic representation). **f)** Set of four environments that impose a single conflict, between and *P*^*10*^ and *P*^*00*^. They cannot cross given the partial order constraint, even though selection favors it. The optimum is then that they are equal (nodes overlap). If *P*^*10*^=*P*^*00*^ increases, *F*^*00*^ increases but at the expense of decreases in *F*^*10*^. The conflict thus identifies a cross-environmental tradeoff. The resulting Pareto front is a one-dimensional line (right, schematic representation), independent of the dimensionality of the fitness space. **g)** Set of four environments that impose two conflicts, and resulting two-dimensional Pareto surface (right, schematic representation).

However, not all environmental switches impose such order. Specifically, it contrasts with situations a switch from an environment *S* = (0,0) to an environment *S* = (1,1), where both *s*_*1*_ and *s*_*2*_ increase. For a single genotype, the order in the two corresponding phenotypes *P*^*00*^ and *P*^*11*^ may be *P*^*00*^ > *P*^*11*^ as represented in Fig. 1c – this is the case when the suppressing effect of *s*_*1*_ dominates over the activating effect of *s*_*2*_. However, this is not necessarily the case, as the activating effect may dominate for some other genotypes. Thus, even when monotonicity is preserved for individual responses, the order between *P*^*00*^ and *P*^*11*^ is not necessarily a constraint during evolution.

### Partial order constraints and their graph representation

The absence of order between some pairs of phenotypic values seems to imply that phenotypic order will not always lead to the identification of a constraint. However, the notion of *partial* order provides an approach to capture the available information on order, which can be represented in graphs called Hasse diagrams^20^. The nodes of the graph represent the phenotypes *P*(*S*) for different environments *S* (Fig. 1d). The node with the highest value of *P* is displayed at the top, the one with the lowest value of *P* at the bottom. Any two nodes that exhibit a specific order are connected by vertices. Here the resulting Hasse diagrams is diamond-shaped (Fig. 1d). Mutations affecting characteristics like activation and repression strength can move nodes up and down, but cannot alter the connectivity or topology of the graph. The graph is a *partial order* graph, as it defines both the order and lack of order that is present.

Under a given evolutionary pressure, the partial order graphs can be reduced to simpler graphs that represent the optimal values for the phenotypes *P*(*S*). For instance, consider the four environments to alternate in time, with low *P* favored only in *S* = (0,1), and high *P* in the other environments (see up or down arrows in Fig. 1e, left). In this problem, all four nodes can optimize to the minimum and maximum *P* values, without encountering conflicts between the posed selective objectives. The solution to this problem (Fig. 1e, middle) is that *P*^*00*^, *P*^*01*^ and *P*^*11*^ take the maximum possible value, and *P*^*10*^ the minimum. This single phenotype determines a single point in fitness space (Fig. 1e, right).

This situation differs from conditions that favor a low *P* only in *S* = (0,0) (Fig. 1f, left), as the latter now conflicts with the selection objective of node *S* = (0,1) (Fig. 1e, middle). The order constraint *P*^*00*^ > *P*^*01*^, here means that the nodes cannot ‘cross’ (as this would result in *P*^*00*^ < *P*^*01*^), and hence not all objectives can be fulfilled. Here, it is optimal to merge the *P*^*00*^ and *P*^*01*^ nodes, meaning *P*^*00*^ = *P*^*01*^. Hence, the system can either optimize *P*^*01*^ (by increasing *P*^*00*^ = *P*^*01*^), or optimize *P*^*00*^ (by decreasing *P*^*00*^ = *P*^*01*^), but not both. In other words, in the absence of the repressing signal (*s*_*1*_=0), the network cannot always produce the optimal phenotype (for both values of the activating signal: *s*_*2*_=0 and *s*_*2*_=1).

### Conflicting objectives, trade-offs, and Pareto fronts

The above conflicts in objectives reflect a trade-off: the fitness in one environment can at some point only increase further by changes that at the same time decrease fitness in another environment. The set of such best possible values of *P* are referred to as the Pareto front^21^. One may note that a certain variable environment does favor a specific location on this Pareto front: the full Pareto front indicates all potential optimal solutions – rather than the single optimum resulting from a specific variable environment. We have just described how, in the presence of a single conflict, the optimum then becomes a line instead of a point in fitness space. Note that the specific curvature of the line depends on the non-linearity of the relation between expression and fitness in each environment (Fig. 1f, right). In the same way, the Pareto front is a surface for two conflicts (Fig. 1g) and so on. We have previously shown theoretically that this approach allows to reduce a large set of potential constraints into a Pareto optimal set of much smaller dimension, starting from arbitrary numbers of signals and monotone responses (Supplementary Fig. 1)^17^. The resolution algorithm developed for this purpose processes the Hasse diagram in order to identify conflicting regulatory objectives, the number of which predicts the dimensionality of the Pareto front (see Supplementary Information).

### Experimental evolution in different environments

We tested these predictions by random mutagenesis of a network, and competitive selection in different consecutive environments that correspond to a variety of objectives (Fig. 2). We engineered a genetic network responding to the inducers doxycycline (dox) and isopropyl-β-D-galactopyranoside (IPTG), which define *s*_*1*_ and *s*_*2*_ respectively (Fig. 2a; Methods; Supplementary Table 1). The network controlled a selection cassette^22^, whose expression level defined *P*. In this manner, the network output *P* was coupled to the growth rate (*F*), on which selection acts (Fig. 2a).

**Table 1.** Fitted values of dissociation constants in a biochemical model of the regulatory networks (See Methods): k_dox_ quantifies the response of the TetR protein to dox, K_tetR_ the binding of TetR to its target promoter (unitless because normalized by constitutive TetR concentration), k_IPTG_ quantifies the response of the LacI protein to IPTG, and K_lacI_ the binding of LacI to its target promoter (unitless because normalized by maximum LacI concentration). For the network as constructed, the fit was done with a TetR induction response; for the intermediates after the first and second rounds, the fit was performed with a TetR co-repression response (Methods).

| Construct | Figure | $k_{\text{dox}}$<br>(ng/mL) | $K_{\text{TetR}}$ | $k_{\text{IPTG}}$ (mM) | $K_{\text{LacI}}$ |
| --- | --- | --- | --- | --- | --- |
| pNetwork-WT | grey dot Fig. 3b | $32 \pm 6$ | $0.011 \pm 0.004$ | $16 \pm 1$ | $0.015 \pm 0.005$ |
| pNetwork-M2 | red dot Fig. 3c | $25 \pm 10$ | $0.05 \pm 0.01$ | $2.5 \pm 0.5$ | $0.04 \pm 0.01$ |
| pNetwork-M3 | orange dot Fig. 3c | $0.23 \pm 0.07$ | $3.3 \pm 0.17$ | $2.5 \pm 0.5$ | $0.04 \pm 0.01$ |

**Figure 2.**
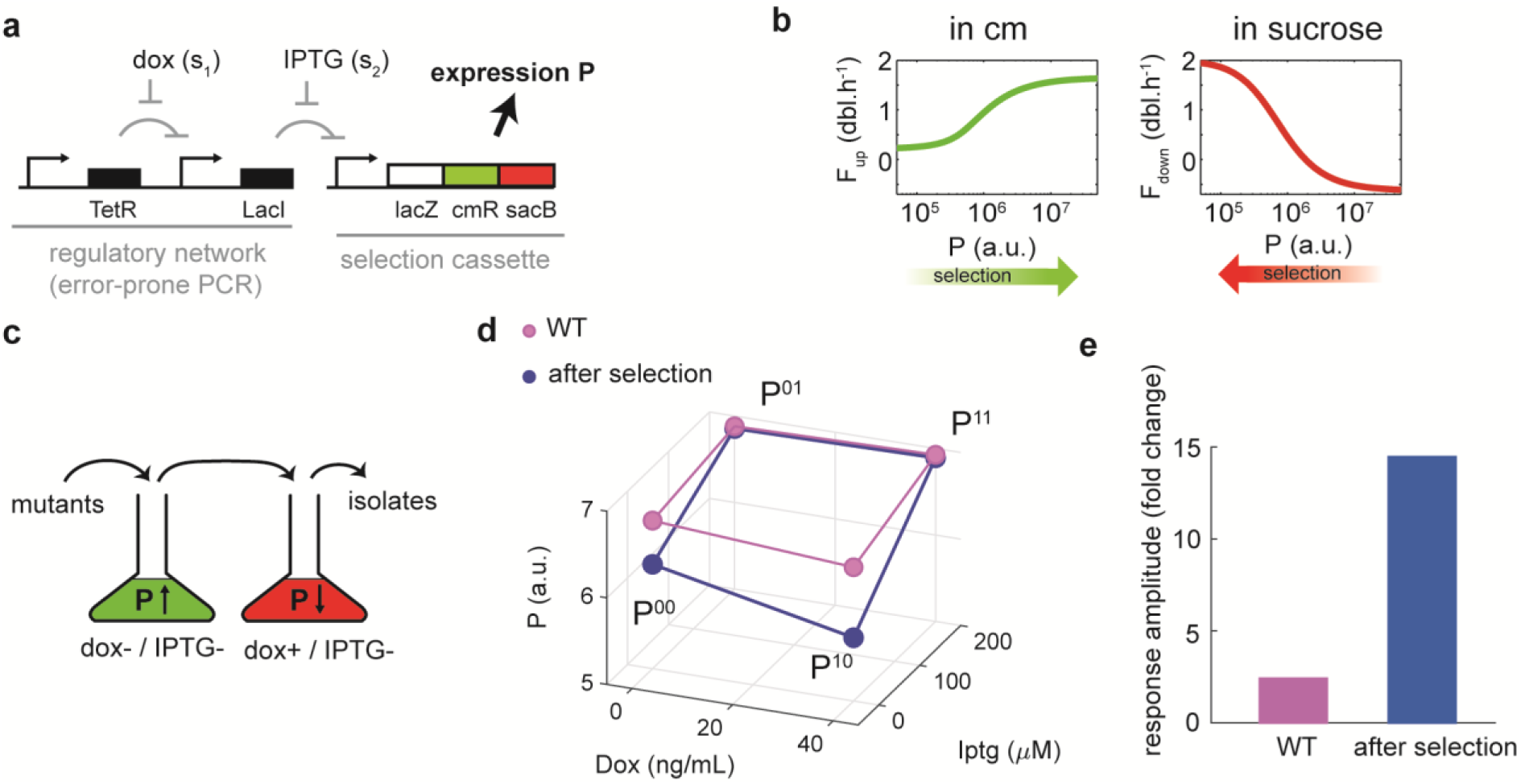
Experimental evolution in multiple consecutive environments. **a)** Engineered genetic network and selection cassette. Expression of selection cassette (*P*) is controlled by two environmental signals dox (*s*_*1*_) and IPTG (*s*_*2*_). The genetic network (black) is mutated using error-prone PCR, yielding a mixed population of different network variants. The selection cassette^22^ contains beta-galactosidase fragment LacZ to measure expression levels, the chloramphenicol resistance gene *cmR* to select for increasing *P*. The levansucrase gene *sacB* confers toxicity in sucrose media. **b)** Quantification of selection in two types of environment (from ref. ^22^). Measured growth rates as function of *P*. For this quantification purpose, *P* is here varied by induction, rather than by mutations^22^. In a chloramphenicol-containing medium (left), cells with high *P* grow faster and hence are favored, due to their high levels of resistance. In a sucrose-containing medium, cells with lower *P* grow faster and hence are favored, due to reduced toxicity. **c)** Illustration of the experimental evolution protocol (see Methods). A mutant population is successively grown in two indicated environments, which couples the input signals to a selection for *P*. **d)** Measured values for the initial network (‘WT’, purple), and after selection (blue) to lower *P*^*10*^, while at the same time preventing TetR knockouts to occur by selecting for high *P*^*00*^. Consistently, *P*^*10*^ decreased, and sensitivity to dox is retained. **e)** Measured increase in the dynamic range, as the fold change between highest and lowest *P* value.

Selection was designed to work as follows. To select for increased *P*, a chloramphenicol-containing medium was used, resulting in a growth rate *F*_*up*_. Increased expression of chloramphenicol acetyltransferase (cmR) within the cassette provides resistance and hence faster growth. The relation between *P* and *F*_*up*_ can be quantified^22^ (Fig. 2b, left), which shows that *F*_*up*_ is close to zero when *P* is low, and increases to about 1.7 doublings per hour (db/hr) when *P* is high. To select for decreased *P*, a sucrose-containing medium was used, resulting in a growth rate *F*_*down*_. Because the polymerization of sucrose by the levansucrase enzyme (SacB) is toxic, decreased expression of the cassette then produces faster growth. Quantification of the relation between *P* and *F*_*down*_ indeed shows that *F*_*down*_ is about 2 dbl/hr when *P* is low and decreases to negative values when P is high (Fig. 2b, right)^22^.

The regulatory behavior of the network corresponds to the example of Fig. 1a. Increases in *s*_*1*_ (dox) should relieve repression by the upstream TetR repressor, of the downstream LacI repressor, which in turn represses the network output *P*. On the other hand, increases in *s*_*2*_ (IPTG) should relieve repression of *P*. Thus, *s*_*1*_ should suppress, while *s*_*2*_ should activate *P* (Fig. 1a). Using enhanced yellow fluorescent protein (eYFP) reporter, we verified that this was indeed the case (Fig. 2d). The dynamic range was small however: even the lowest *P* (*P*^*10*^) was near the end of the range of *P* values (2.4-fold change, Fig. 2e). To validate the experimental evolution protocol, we first aimed to select for a lower *P*^*10*^.

The network (i.e. the TetR and LacI coding sequences, their promoters and their operator sites) was randomly mutated by error-prone PCR with on average 3 mutations per gene, and subsequently inserted into a vector containing the selection cassette. This resulting pool of mutant networks was transformed into *E*. *coli* and selected in a medium with chloramphenicol, but without any inducers, which corresponds to an upward pressure for *P*^*00*^. After this the population was transferred to a medium with dox, without IPTG, and containing sucrose (Methods), which corresponds to a downward pressure on *P*^*10*^. This dual selection provided a counter-selection for TetR knock-out mutants, which can be achieved by a wide range of mutations, and indeed would decrease *P*^*10*^, but also fully abolish the response to dox. Finally, isolates of the resulting population were characterized using an eYFP reporter gene for the network output (Fig. 2d). This indicated that *P*^*10*^ had indeed decreased significantly (Fig. 2e, 14-fold change of the response after selection) while the response to dox remained intact.

### Fitness domains constrained by partial order

To test the partial order approach, we focused on the case of conflicting objectives (Fig. 1f). This scenario corresponded to a downward selection of *P*^*00*^, an upward selection of *P*^*10*^, *P*^*01*^, and *P*^*11*^ (Fig. 3a), and hence a conflict between *P*^*00*^ and *P*^*01*^ (Fig. 3a, dashed line). Using the predicted optimum under partial order constraints (Fig. 1f, middle) and the measured *F*_*down*_(*P*) and *F*_*up*_(*P*) (Fig. 2b), we determined the shape of the predicted 1-dimensional Pareto front (Fig. 3b, orange line). We could further determine the accessible 4-dimensional fitness space in the 4 environments (Supplementary Fig. 2), here visualized in 3 dimensions by averaging the fitness of *P*^*10*^ and *P*^*11*^ (Fig. 3b). In this fitness space, each possible network genotype corresponds to a single point. Note that the plotted dots in Fig. 3b are measured experimental phenotypes obtained after the selection protocol described in the next section. In this same space, the Pareto front represents a collection of networks. The two extreme ends of this Pareto line are two archetypal regulatory responses (Fig. 3b), following Shoval *et al*.^23^. These respectively correspond to *P*^*00*^ and *P*^*01*^ being low, while *P*^*10*^ and *P*^*11*^ are both high, and *P*^*00*^, *P*^*10*^, *P*^*01*^, and *P*^*11*^ all high (Fig. 3b). The leveling off of *F*_*down*_(*P*) and *F*_*up*_(*P*) at low and high *P* (Fig. 2b) define the lower and upper limit of the Pareto line (Fig. 3b). The other corners of the fitness boundaries in Fig. 3b correspond to other archetypal regulatory phenotypes that are not optimal under this selective regime (Supplementary Fig. 2). Altogether, the regulatory archetypes and the line which connect them limit a domain of fitness that is accessible under the defined partial order constraints, while the space beyond it is predicted to be inaccessible.

**Figure 3.**
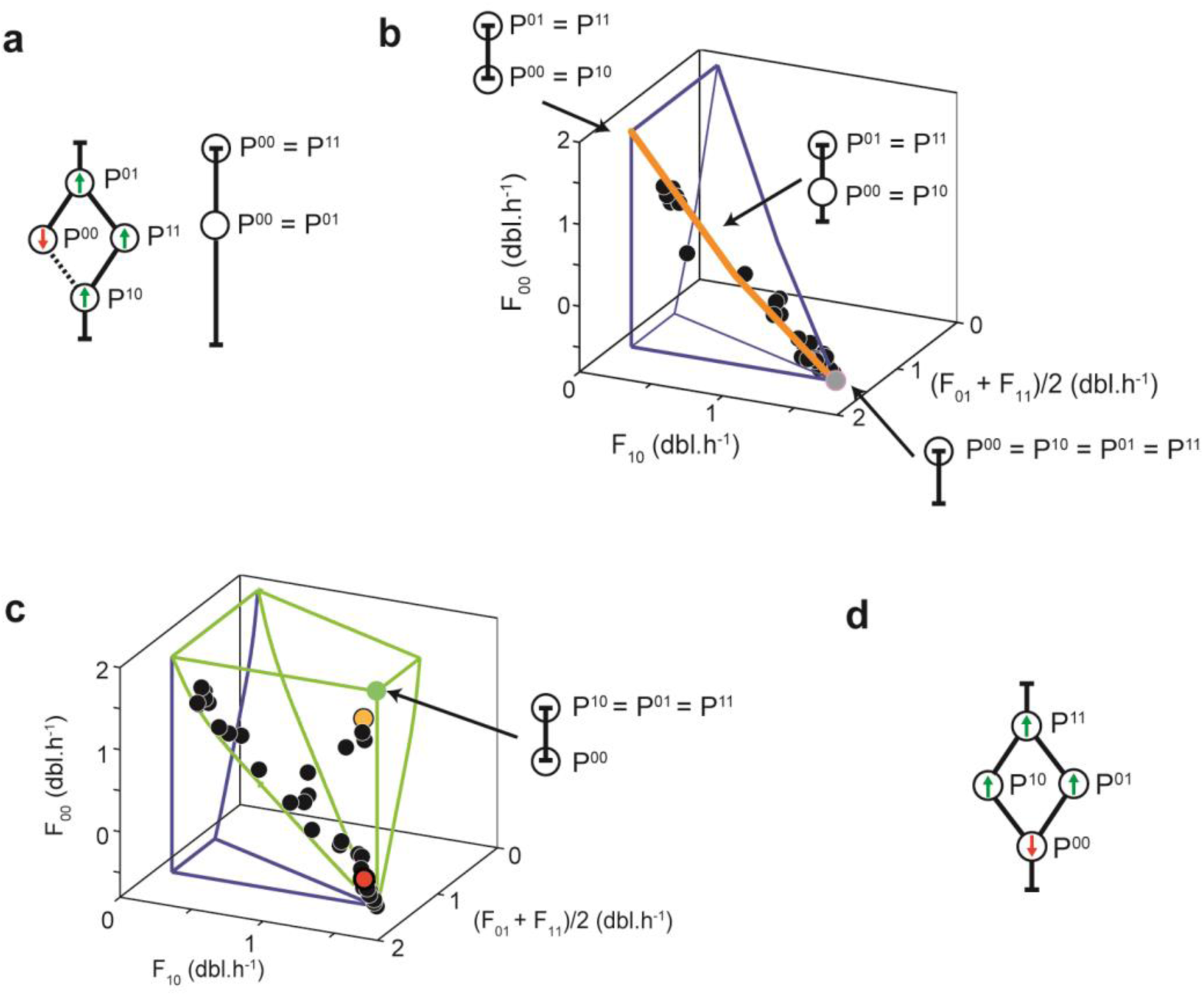
Experimental evolution of conflicting objectives. **a)** Left: Partial order graph with (conflicting) selective pressures indicated by arrows (as in Fig. 1f). Dashed line indicates a single conflict between *P*^*10*^ and *P*^*00*^, which results from incompatibilities between the selective pressures and the partial order graph. Right: Pareto optimal solution. **b)** Predicted accessible fitness domain for four environments and isolates after the first round of evolution. The fitness values of two environments are averaged for visualization purposes. Corners indicate predicted optimal archetypal regulatory phenotypes, connecting edges represent intermediate regulatory phenotypes. Edges are drawn using the indicated changes in the values of *P* (which here all move between the minimum and maximum values), the defined selective pressures (panel a), and the corresponding measured relations between *P* and fitness (Fig. 2b). The orange line is the one-dimensional Pareto-optimal front. The grey dot represents the initial regulatory phenotype and is close to an archetypal phenotype where expression is maximum in all environments. During experimental evolution, a population of cells with randomly mutated networks is exposed to selection in multiple sequential environments. Black dots are isolates of the resulting population and scatter along the predicted Pareto front. **c)** Predicted accessible fitness domains and isolates after the second round of experimental evolution. Some isolates continue to scatter along the Pareto front. Others moved away from it, suggesting the constraint is broken by changes in the partial order. Green lines: predicted accessible space for adjusted partial orderings. Isolates approached the optimum (zero-dimensional Pareto front, green dot). The orange dot represents the most optimal regulatory phenotype measured. The green dot represents the most optimal phenotype reachable in theory. **d)** Adjusted partial order with conflict resolved. *P*^*10*^ and *P*^*00*^ now crossed and swapped places, consistent with the selective pressure (arrows). *P*^*01*^ and *P*^*11*^ also crossed, as dox is now a suppressing signal.

### Experimental evolution with conflicting objectives

The initial network (Fig. 2a) was found to map to a corner of this fitness domain (Fig. 3c, grey dot) and to correspond to an archetypal response. This position is consistent with the empirical observation that *P* was high in all four environments (Fig. 2d). Next, we mutated the network by error-prone PCR and performed selection in different environments, with a protocol similar to the one described in Fig. 2c, but with the selective pressures as indicated in Fig. 3a. We found that the resulting isolated genotypes remained confined within the fitness domain predicted by the partial order. Moreover, they scattered along a line within the multi-dimensional fitness space, which coincided with the predicted Pareto front (Fig. 3b, black dots along the orange line, Supplementary Fig. 3a). These selected genotypes showed increased *F*^*00*^, at the expense of decreased *F*^*01*^, while little affecting *F*^*10*^ and *F*^*11*^ (Fig. 3b), consistent with the single conflict identified between the nodes *P*^*00*^ and *P*^*01*^ (Fig. 3a).

A second round of mutagenesis and selection was performed, using one of the selected isolates as a founder (Fig. 3c, red dot). Consistent with the previous round, many of the progeny that were present in the population after selection in multiple environments, were again found scattered along the 1-demensional Pareto front (Fig. 3c). Some of the progeny however, had moved outside the predicted fitness domain (Fig. 3c). These isolates were scattered along a line that bridged the Pareto front and the theoretical fitness optimum (green dot, Fig. 3c, Supplementary Fig. 3b), where growth rates are as high as they can be in all environments. These findings could, at least in principle, indicate a problem with our predictions, or rather indicate the emergence of genetic variants that resolved the conflict between the *P*^*00*^ and *P*^*01*^ objectives, and hence overcame the partial order constraints.

### Breaking order constraints with changes in network structure

We aimed to assess if the partial order framework was consistent with the movement outside the fitness domain (Fig. 3c). If so, and a conflict was thus resolved, the partial order should have changed from the original to a new one. Originally, the objectives of decreasing *P*^*00*^ while increasing *P*^*01*^ conflicted with the phenotype order *P*^*00*^ > *P*^*01*^ (Fig. 3a). This conflict would be resolved by a changed order: *P*^*01*^ > *P*^*00*^ (Fig. 3d). If this hypothesis is correct, it should be manifested by a corresponding change in the underlying network structure. The predicted new partial order (Fig. 3d) can be the result of a several potential changes in the network, representing the various ways in which the signal s_1_ (dox) would no longer suppress *P* but rather activate it (Fig. 4a, from top to bottom): 1) the upstream TetR repressor becomes an activator, 2) the downstream LacI becomes sensitive to dox, 3) dox becomes a co-repressor of TetR, and 4) the upstream TetR becomes a direct repressor of the output, like LacI. In these solutions, not only the order between *P*^*00*^ and *P*^*01*^ changes, but also the order *P*^*10*^ > *P*^*11*^ reverses to *P*^*11*^ > *P*^*10*^ as in Fig. 3d. For networks with both order reversals, the corresponding fitness domain is directly adjacent to the original one, and indeed contains the network variants that moved away from the initial Pareto constraint (Fig. 3c, green lines).

**Figure 4.**
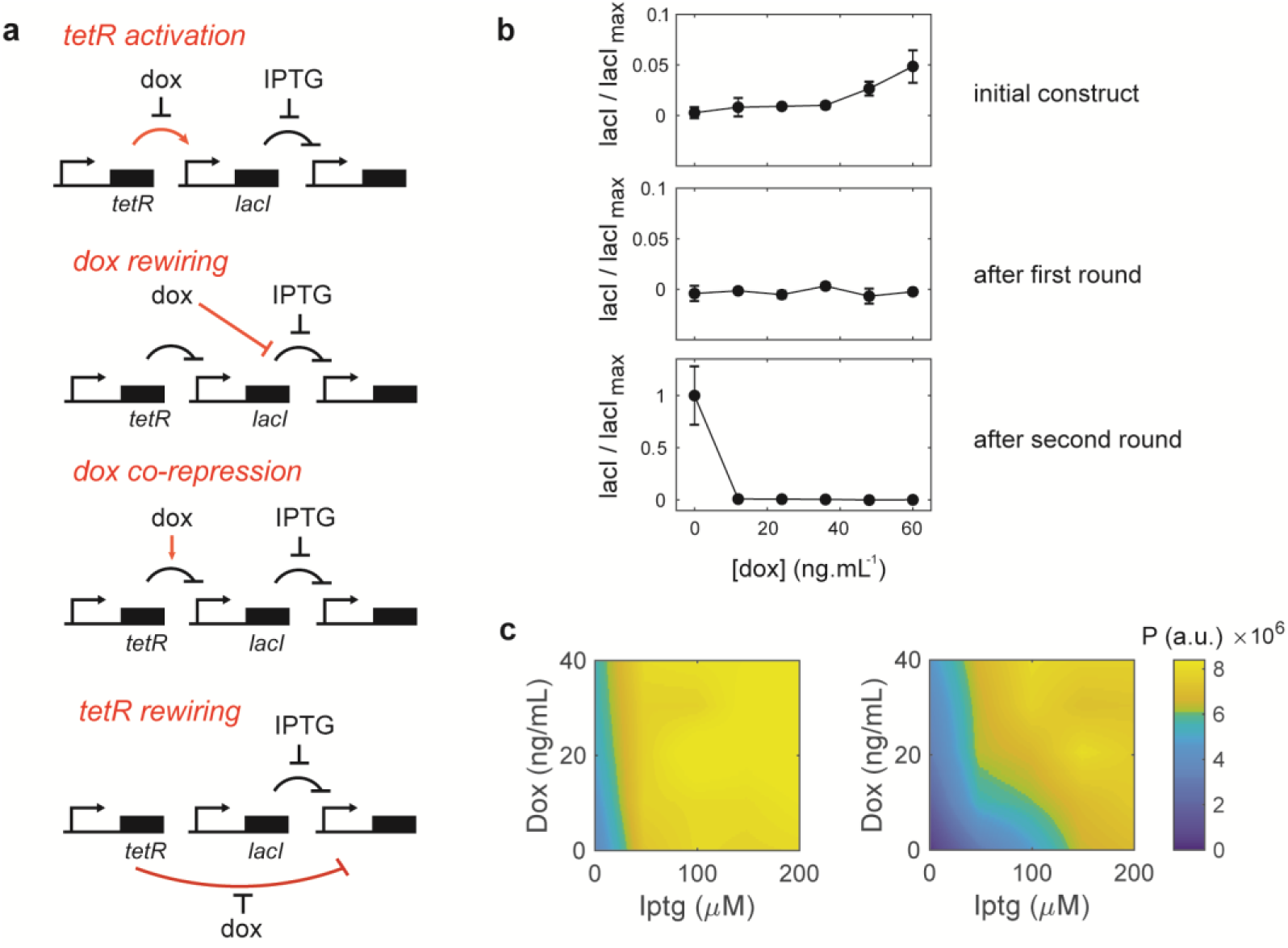
Breaking an order constraint with network innovations. **a)** Four possible network changes that all correspond to the adjusted partial order, in which the conflict (Fig. 3a, dashed line) resolved (Fig. 3d). In all cases, dox now activates *P* whereas it suppressed *P* originally (Fig. 2a). **b)** LacI expression level vs dox concentration for different TetR variants. LacI expression level is normalized by the highest measured value measured across all experiments (no dox, lowest panel). LacI expression is measured by fusing it with a fluorescent marker. Top: data for network as constructed. Dox relieves repression of LacI by TetR, and hence increases LacI expression. Middle: LacI expression after first round, of isolate that was transferred to the second round (Fig. 3d, red dot). Expression is now insensitive to dox, within detection limit. Bottom: LacI expression after the second round, of most optimal isolate (Fig. 3c, orange dot). Dox now decreases LacI expression. **c)** Measured *P* as a function of dox and IPTG, for the isolate transferred from round 1 (panel c, middle), and the most optimal isolate (panel c, bottom). These data are consistent with dox having become a co-repressor of TetR (panel a, bottom).

In order to distinguish between the different options (Fig. 4a), we sequenced and characterized the responses of the most optimal network variants after round 1 and 2 (Supplementary Fig. 4, Supplementary Table 2). The sequences revealed point mutations in the evolved *tetR* and *lacI* coding sequences, but not in the regions controlling DNA or ligand binding, nor in non-coding regulatory regions. Solutions involving altered ligand binding, which could make LacI sensitive to dox (Fig. 4a, second from the top), and altered TetR binding sites on the DNA, which could enable an activator TetR variant (Fig. 4a, top), or allow TetR to repress the output directly (Fig. 4a, bottom), thus seem unlikely. To further test this hypothesis, we functionally characterized the most optimal TetR variants at the end of the first and second rounds, by measuring the expression of a fluorescent protein that they controlled directly (Methods). These data showed that fluorescent protein expression increased with increasing dox for the wild-type TetR, as expected for an induced repressor (Fig. 4b). Expression became low and insensitive to dox after the first round. However, after the second round, expression was high for low dox and then decreased with increasing dox (Fig. 4b). These data suggest that dox has become a co-repressor of TetR (Fig. 4a, third from the top). The expression *P* as a function of dox and IPTG concentrations (Fig. 4c) was indeed improved; from a nearly insensitive response to dox before the second round of selection (Fig. 4c, top), to a regulatory phenotype (Fig. 4c, bottom) in which *P*^*00*^ is minimized and *P*^*10*^, *P*^*10*^, *P*^*11*^ are maximized. In fact, the observed mutation R49G (Supplementary Table 2) is consistent with a TetR inversion^24^. Fitting these data to a cascade model of biochemical rate constants suggests that the dissociation constants of both transcription factors have changed across the rounds of evolution (Table 1, Supplementary Fig. 5). Overall, we found that a molecular innovation resolved the conflict and allowed access into a new region of fitness space.

## Discussion

Pareto fronts have long been established as a powerful concept in disciplines ranging from economy to instrument design^25–27^. They allow one to consider possible solutions and their limits when pursuing multiple objectives. In biology, Pareto fronts are observed when mapping collections of species within multi-dimensional phenotype spaces^28,29^. It has been proposed that Pareto fronts can be detected by interpolating highly specialized phenotypes (also called archetypes)^23^. They thus provide insight in constraints arising from functional trade-offs in evolutionary adaptation. At the same time, it is a major challenge to mechanistically understand and predict such constraints^3,5,10,30^, in addition to observing them. Even framing the problem is not straightforward: to identify what aspect could be predictable and what not, and which information about the evolving system is then required, whether it be at the genetic, phenotypic, or fitness level.

Here, we addressed these issues by developing a framework to predict constraints of networks that integrate multiple signals in monotonic fashion. We found that the notion of partial order identifies such constraints. Specifically, it defines the evolutionary limits of a network, in which functional properties such as transcription factor binding affinities can be altered, but their activating or repressive nature and the overall network topology remains unchanged. The partial order captures the limited amount of information that is needed, namely whether input signals activate or suppress the phenotype in question. Notably, not needed are typically poorly understood details like the actual topology, the number of regulatory proteins, how they function, or which mutations affect their function and how. Owing to its foundations in graph theory, the approach is well suited for more complex environments and regulatory objectives, and indeed can reveal the minimal core underlying conflicts between these objectives (for example see Supplementary Fig. 1)^17^.

The partial order framework may be used to predict the space of accessible phenotypes, provide the dimensionality of the Pareto-optimal front, its shape, identify extremal regulatory phenotypes (regulatory archetypes), and allow more targeted network engineering approaches. Interestingly, it also directly provides predictions on the dimensionality of the Pareto-optimal front, which equals the number of conflicts between (regulatory) objectives in different environments. This dimensionality of the Pareto front relates to diversity, as a zero-dimensional front indicates a single optimal phenotype, while additional dimensions indicate that diverse phenotypes can be equally optimal. Conceptually, one may see the partial order analysis to apply to regulatory networks in a similar fashion that Flux Balance Analysis^31^ applies to metabolic networks, where the mere knowledge of a graph structure constraints accessible fluxes, and optimality is used to predict evolutionary outcomes. We note that the approach is not valid for non-monotonic responses to signals, when unknown signals vary jointly with considered signals, and when dynamical features such as oscillations are central to function and selective advantage. On the other hand, many regulatory responses are monotonic^19^. In addition, it has been shown that any biological regulatory networks can be decomposed into monotonic modules^32^, which may allow further generalization.

The experiments presented here provided a direct test of these concepts, and illustrate which types of functional changes can modify the phenotype order. Experimental evolution in multiple environments revealed two modes of adaptation. In the first mode, solutions that emerge are those that obey the partial order constraints defined by the founding genotype, and are enriched at the predicted Pareto-optimal front. The second mode involves a type of mutations that are rarer: those that confer functional innovations which are able to alter the partial order, and hence allow escape from these constraints. The findings thus identify qualitatively distinct evolutionary stages in regulatory strategies.

The ability to define evolutionary constraints of regulatory phenotypes, as we have pursued here, will be central to arrive at a mechanistic understanding of evolution in complex niches. It can provide hypothesis on the compatibility of different regulatory objectives, or lack thereof, and on their evolutionary accessibility, as also illustrated by our data. Such regulatory limitations, and associated tradeoffs when occupying broad spectra of environmental conditions, can promote niche exclusion in the context of competition^33^, and hence play a role in species diversity and coexistence.

## Materials and Methods

### Constructs

We modified a regulatory circuit in which the selection operon consisting of *lacZα, cmR*, and *sacB* genes driven by the promoter *P*_*trc*_ is under control of the LacI transcriptional repressor^22^. Expression of LacI is under the control of TetR repressor via promoter *P*_*LtetO1*_. TetR itself is constitutively expressed via promoter *P*_*N25*_^22^. This network harboring plasmid contains a kanamycin-resistance gene and a medium copy *p15A* origin of replication. Materials are available upon request.

To measure the output from the network variants, various reporter constructs were used (Supplementary Table 1). The LacZ based assays utilized either constitutively expressed *lacZω* fragment via *P*_*lacI*_^*Q*^ to measure in *cis*, that constitute a functional LacZ together with *lacZα* encoded by the selection operon of the network, or the full version of *lacZ* under the promoter *P*_*lac*_ to measure in *trans*^22^. Whilst the fluorescent protein based readout assays utilized the plasmid encoding either *lacI-mCherry* under the promoter *P*_*LtetO1*_ or *eYFP* under the promoter *P*_*trc*_ ^16^. This reporter plasmid backbone contains an ampicillin-resistance gene and a medium copy *colE1-rop* origin of replication, which is compatible to co-reside with the network harboring plasmid in the same cell (Supplementary Table 1).

### Mutagenesis

The mutations were introduced into the regulatory network sequence spanning *tetR* to *lacI* including the promoters of respective genes by error-prone PCR (Stratagene Genemorph II Random Mutagenesis kit). The mutated PCR amplicons digested with DraIII (NEB), ligated into the vector backbone containing the intact selection cassette, and transformed into *E*. *coli MC1061* strain by electroporation (Avidity EVB100). Routinely we obtained a pool size of half a million to ten billion. To determine the mutation rate, DNA isolated from randomly picked transformants was Sanger sequenced, revealing an average mutation rate of 3.0 mutations per kb (n=9).

### Selection

Cells harboring mutant networks were grown at 37 °C with vigorous shaking in 20-40 mL of EZ Rich Defined medium (Teknova, Cat. M2105) supplemented with 0.2% glucose as a carbon source, 1 mM thiamine hydrochloride and 50 µg/ml kanamycin. The small-molecule inducers doxycycline or isopropyl-β-D-galactopyranoside (IPTG) were added 3 hours prior to the beginning of selection. After this pre-selection growth phase, chloramphenicol (40 µg/ml) or sucrose (0.25% w/w) were added for selection and cells were cultured for an additional 6 hours. This duration of the selection growth phase was chosen to obtain significant enrichment factors (of up to 10^4^), while still maintaining the diversity of the population. During selection the optical density of the culture was monitored at regular intervals and diluted 500 times in pre-warmed medium whenever the optical density (OD) at 550 nm reached 0.1 value.

### Measurement of network responses

The output of an isolated network variant was measured by co-transforming with a suitable reporter encoding plasmid (Supplementary Table 1).

For LacZ based assays, 200 µl cultures were grown at 37 °C in EZ Rich Defined medium (Teknova, cat. M2105) with glucose as a carbon source and supplemented with 1 mM thiamine hydrochloride and the appropriate antibiotics, in a 96-well optical-bottom black color micro-titer plate (NUNC, Cat. 165305), using Wallac Perkin Elmer Victor3 plate reader. The OD at 600 nm was recorded at regular intervals of 4 minutes, and evaporation of the cultures were contracted by adding 9 µl of sterile water per well at an interval of 29 minutes. When most of the cultures were grown to 0.05 to 0.2 OD range, cells were fixed by adding 20 µl of fixation solution, which was freshly constituted before use by mixing 109 µM fluorescein-di-β-D-galactopyranoside (FDG, MarkerGene, cat. M0250) substrate, 0.15% formaldehyde and 0.04% DMSO in sterile water. The development of the fluorescence from LacZ activity was measured at regular interval of 8 minutes by excitation at 480 nm and emission at 535 nm, in parallel to the OD_600_ measurement. This data was analyzed as described previously^22^.

For fluorescent protein based assays, the cultures were grown in EZ Rich Defined medium early-exponential growth phase, and then diluted into a final OD_550_ of 1×10^−4^ and transferred to a 96-well optical-bottom black color micro-titer plate (NUNC, Cat. 165305) in a total volume of 200 μL per well. The OD_550_ and fluorescence intensities from two distinct fluorescent reporter proteins (mCherry (excitation 580/20, emission 632/45) and eYFP (excitation 500/20, emission 535/25) were monitored in a Wallac Perkin Elmer Victor3 plate reader at regular intervals at 37 °C. The instrument was shaking (double orbital) and replenishing 9 μL of sterile water per well every 27 minutes. This data was analyzed as described previously^16^.

### Fitness computation

The fitness as a function of the expression level *P* was previously modeled and fitted^22^ and take the form:

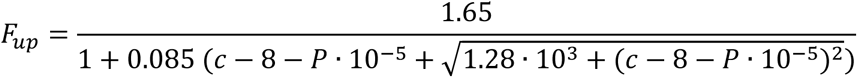

where *F*_*up*_ is the increasing fitness response as a function of increasing expression levels *P*, in the presence of a chloramphenicol concentration *c* = 40 µg/ml in the medium, and

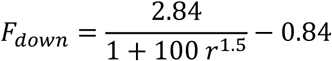

with

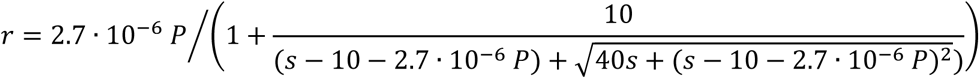

where *F*_*down*_ is the decreasing fitness response to increasing expression levels *P* in the presence of a sucrose concentration *s* = 0.25% (weight fraction) in the medium. Expression levels of the mutants of Figure 3 are reported in the Supplementary Information.

### Estimation of network parameters

The binding constants of Table 1 were estimated by fitting the responses of the separate components (Supplementary Figure 5) assuming constant constitutive expression of TetR and using the following forms:

i. TetR induction by dox:

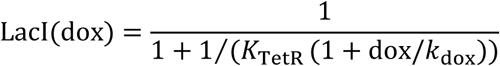
ii. TetR co-repression by dox:

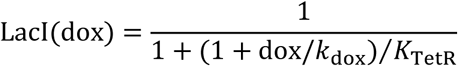
iii. YFP output as a function LacI induction by IPTG:

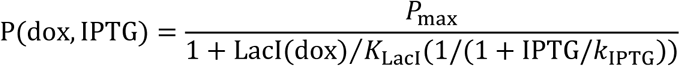

where *K*_TetR_ is expressed in units of constitutively expressed TetR concentration, LacI and *K*_LacI_ are both normalized to maximum LacI expression. Fitted curves shown in Supplementary Figure 5. For each regulatory network, the four dissociation constants where fitted to 31 experimentally measured data points which were measured in duplicate.

## Acknowledgements

Work in the S. J. T. group is part by the Netherlands Organization for Scientific Research (NWO).

## Author contributions

The research was conceived by MK, PN, FJP and SJT. The experiments were performed by MK and FJP. Data was analyzed by MK, PN, FJP and SJT. Theory was developed by PN and SJT. Manuscript was written by PN, MK, FJP, and SJT.

