## Supplementary Material for "Evolutionary constraints in regulatory networks defined by partial order between phenotypes"

Kogenaru et al.

**Mathematical formulation of Pareto optima in monotone regulatory phenotypes**

We consider the environments $E^{(n)}$ (n = 1, ..., N), each corresponding a vector $S^{(n)}$ of inputs signals $(s_1^{(n)},\dots,s_K^{(n)})$ and a monotone regulatory network which transforms each input vector $S^{(n)}$ into a certain level $P^{(n)}$ of the phenotype. Given the monotonicity of the network, the output $P^{(n)}$ is an increasing (resp. decreasing) function of $s_k^{(n)}$for the indexes k in the subset $I^+$ (resp. $I^-$). The natural partial order on $\mathbb{R}^K$ induces a partial order "≥ " between the $P^{(n)}$ values defined as follows: $P^{(u)} \geq P^{(v)}$ if and only if for every k in $I^-$, we have $s_k^{(u)} \leq s_k^{(u)}$, and for every k in $I^+$, we have $s_k^{(u)} \geq s_k^{(u)}$. Additionally, depending on the phenotype being beneficial or deleterious, we consider a fitness optimization objective being respectively the maximization or minimization of $P^{(n)}$.

We have devised a graph algorithm which allows to compute efficiently (in polynomial time) an exact closed form of the Pareto front of this optimization problem under partial order constraints as reported in [S1]. The steps of the algorithm are exemplified in Supplementary Fig. 1. The problem is represented as a directed graph, where each vertex u corresponds to the phenotype level $P^{(u)}$ in the environment $E^{(u)}$, and an edge between from vertex u to vertex v indicates $P^{(u)} \geq P^{(v)}$. To each vertex, we attribute one of the four following states:

1. "descending" if $P^{(u)}$ should be minimized;
2. "ascending" if $P^{(u)}$ should be maximized;
3. "trade-off" if the vertex results from the fusion of a descending and an ascending vertices;
4. "bound" if the vertex results from the fusion of a vertex representing the minimum or the maximum possible value of $P^{(u)}$.

The solution to the problem is found by an algorithm applied to the graph of the partial order between the $P^{(n)}$ values, where the vertex states are attributed according to fitness objectives. At each step of the algorithm, a pair of vertices {U,V} is fused (corresponding to imposing $P^{(u)} = P^{(v)}$), according to the rules described below, applied recursively to the graph, until there is no ascending or descending vertex left:

- Perform a transitive reduction of the graph.
- Find an ascending (or resp. descending) vertex V which is pointed to (resp. which points to) no other vertex of the same state.
- For each vertex U which points to (resp. is pointed by) V, create a new graph resulting from the fusion between U and V, the state of this vertex being "bound" if U is a "bound", or " trade-off" in any other case.

We have shown in [S1] that the terminal graphs of the recursion are exact and minimal parameterizations of each face of the Pareto front. They consists of convex sub-spaces of $\mathbb{R}^N$ defined by the partial order relations (given by a terminal graph) between sets of variables all equal to each other (the $P^{(u)}$ values which are grouped in a same vertex). The union of all these subspaces is the

Pareto front. The number of vertices of each terminal graph provides the dimension of the corresponding face.

### Application to signal integration networks

The full envelope of the accessible fitness, as depicted in main text Figure 3, has been obtained by computing all the extremal regulatory phenotypes compatible with the partial order (Supplementary Figure 2). Given that the accessible domain of regulatory phenotypes is convex, the envelope of the domains can be computed by interpolating their extremal phenotypes. The envelope in the space of fitness values is then the image $F(P^{(1)},\ldots,P^{(N)}) = (F_1(P^{(1)}),\ldots,F_N(P^{(N)}))$, where each $F_n$ is either the $F^+$ or the $F^-$ function of Fig. 2b depending on whether the environment $E^{(n)}$ contains respectively chloramphenicol or sucrose.

| Construct/strain | Marker | Comment |
|---|---|---|
| | | **Networks** |
| pNetwork-WT | Kanamycin | Wild-type (WT) network (purple Fig. 2d, grey dot Fig. 3b) |
| pNetwork-M1 | Kanamycin | Mutant network (blue Fig. 2d) |
| pNetwork-M2 | Kanamycin | Mutant network (red dot Fig. 3c) |
| pNetwork-M3 | Kanamycin | Mutant network (orange dot in Fig. 3d) |
| | | **Reporters** |
| p$P_{lacI^{Q}}$-*lacZω* | Ampicillin | Used to measure the networks output in *cis* from the populations that survived the selection by a LacZ assay |
| p$P_{lac}$-*lacZ* | Ampicillin | Used to measure the networks output in *trans* from the populations that survived the selection by a LacZ assay |
| p$P_{trc}$-*eYFP* | Ampicillin | Used to measure the networks output of the chosen isolates by an orthogonal fluorescence assay |
| p$P_{LtetO1}$-*lacI-mCherry* | Ampicillin | Used to measure only the TetR output in the WT and mutant networks |

**Supplementary Table 1: Constructs used in this study.** The network encoding plasmid vectors were co-transformed with one of the reporter vectors to measure the output of the networks.

| Mutants | $P_{N25}$ | *tetR*/protein | *lacI*/protein |
|---|---|---|---|
| pNetwork-M1, purple Fig. 2d | - | G128C/Trp43Ser | G531A/synonymous |
| | - | C132T/synonymous | C1001T/Thr334Met |
| | - | C149G/Ala50Gly | - |
| | - | G214T/Gly72Trp | - |
| | - | C568A/Leu190Ile | - |
| pNetwork-M2, red dot Fig. 3c | - | C132T/synonymous | G285A/synonymous |
| | - | C145G/Arg49Gly | G325T/Ala109Ser |
| | - | C271Δ/Leu91Δ | C900T/synonymous |
| | - | T272Δ/Leu91Δ | - |
| | - | A273Δl/Leu91Δ | - |
| pNetwork-M3 orange dot Fig. 3c | A-36G | C132T/synonymous | G285A/synonymous |
| | - | C145G/Arg49Gly | G325T/Ala109Ser |
| | - | C271Δ/Leu91Δ | C900T/synonymous |
| | - | T272Δ/Leu91Δ | - |
| | - | A273Δ/Leu91Δ | - |
| | - | T355A/Phe119Ile | - |
| | - | G549T/Glu183Asp | - |

**Supplementary Table 2**: **Genotypes of the studied networks.** The network encoding plasmid vectors were isolated from the *E. coli MC1061* strain and subjected to Sanger sequencing. Only one mutations occurred in the promoter regions ($P_{N25}$).

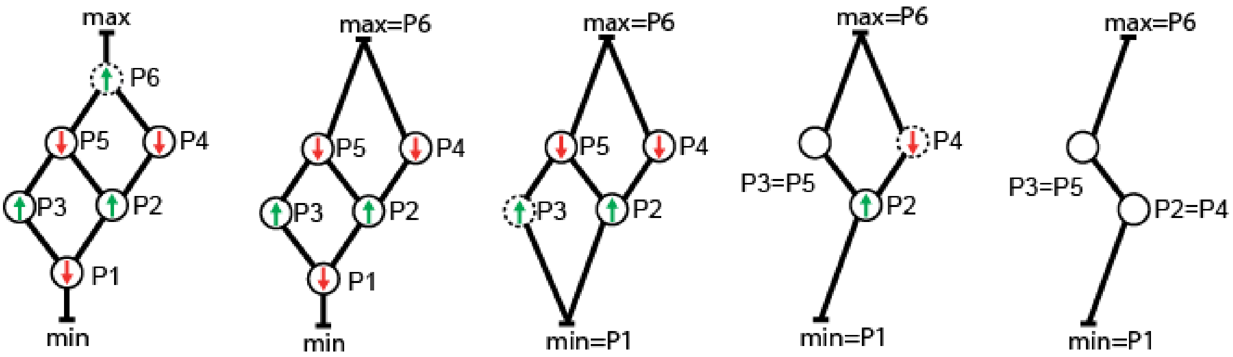


**Supplementary Fig 1. Pareto front parameterization by graph contractions**. To illustrate the generality of the graphical simplification of a partial order with fitness objectives, we consider here an example problem with 6 rather than 4 different phenotypes. The initial problem is represented on the left, leads to the solution on the right after a number of edge contraction as described in the Supplementary Information text. At each step, the node to be merged with another node has a dotted contour. In this example, we see that, in the initial problem, some nodes are confronted to a trade-off with multiple other nodes (ex: P2 is in conflict with P5 and P4). The algorithm allows to minimize the number of such alternative node fusions and find a minimal but exhaustive representation of the full Pareto front. Here, ultimately, the front has dimension 2 despite the fact that 3 pairs of nodes are in conflict in the initial problem (P3-P5, P2-P5, and P2-P4). The analysis thus shows that the P2-P5 conflict is embedded in the other conflict resolutions (consistently, the situation P2=P5 is represented in the terminal graph on the right, as the two middle nodes are allowed to collapse, but not cross).

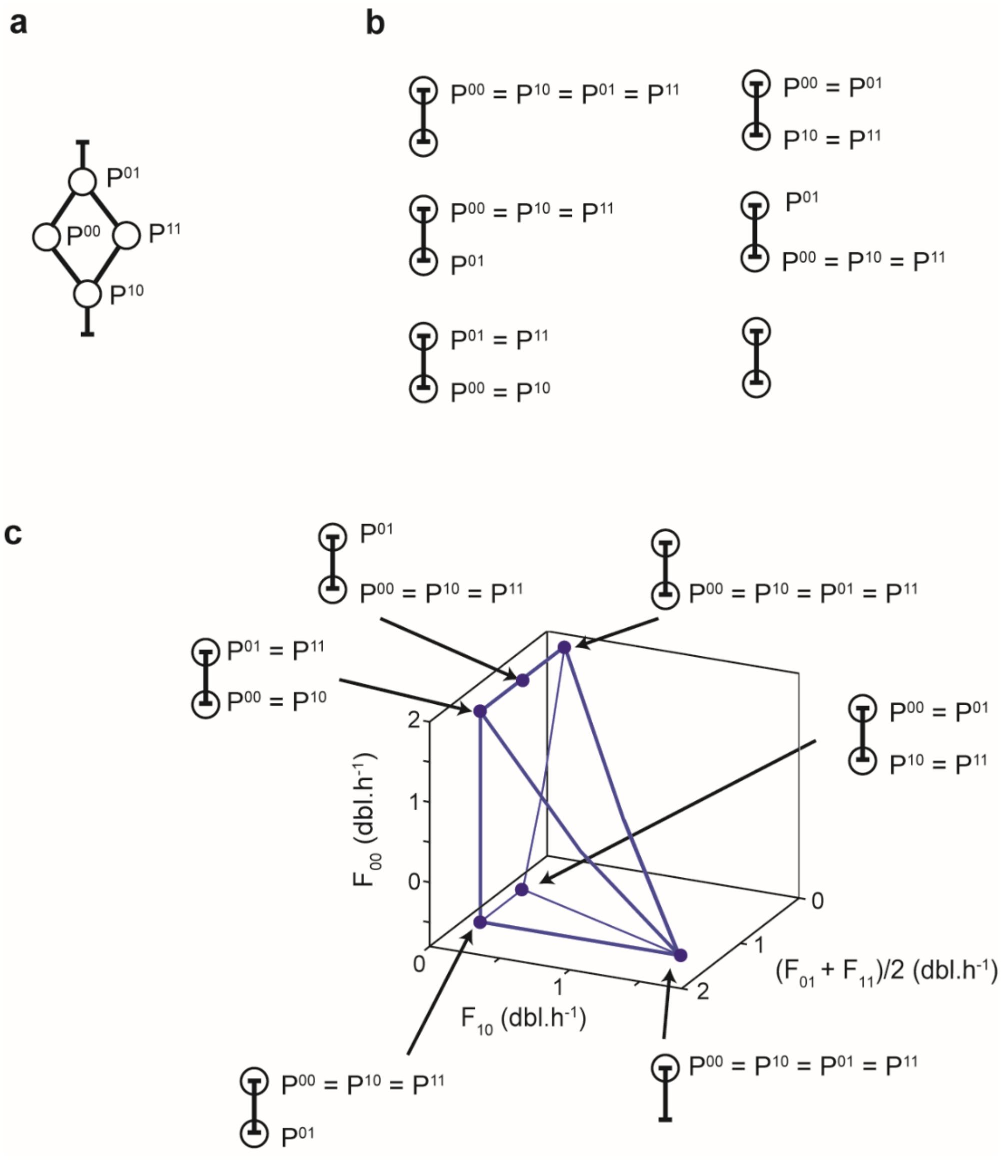


**Supplementary Fig 2: Determination of fitness domain from extremal regulatory phenotypes.** a) Partial order constraints. b) Extremal regulatory phenotypes compatible with the partial order, defining the corners ('regulatory archetypes') of the accessible phenotype space. c) Mapping and projection of the accessible fitness space, obtained by taking the image of the phenotype domain by the fitness functions defined in Fig. 2b and the selective pressures defined in Fig. 3a.

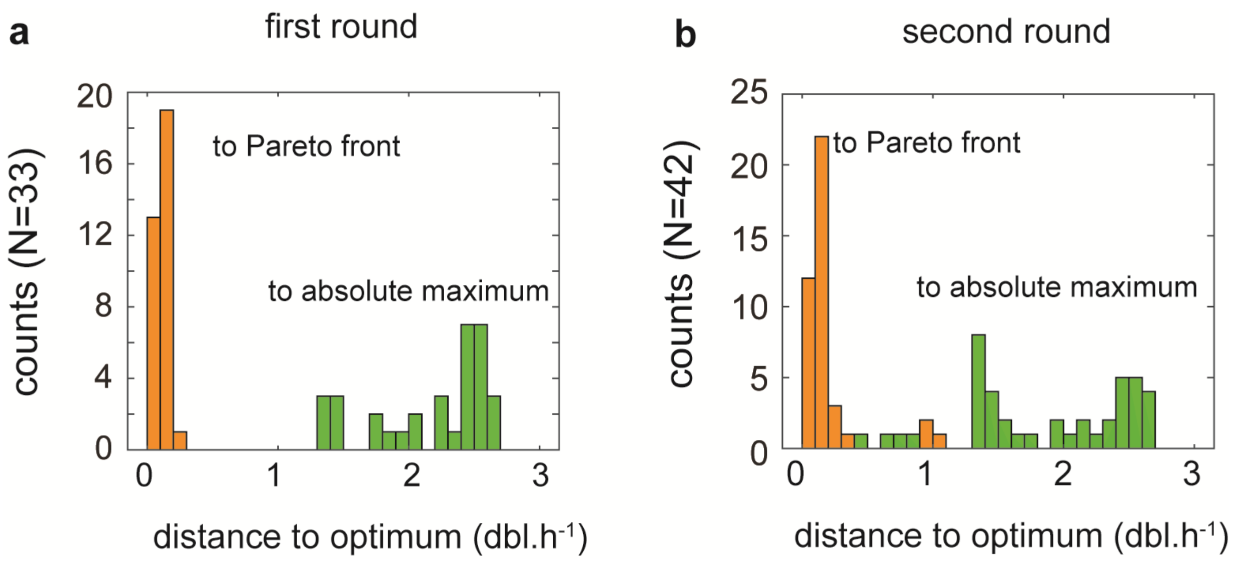


**Supplementary Fig 3: Euclidian distance to optima in the 4 dimensional fitness space. a)** Distribution of Euclidian distance of the mutants after the first round of selection (black dots of Fig. 3b). Orange bars: distance of the mutants to the Pareto front (orange line in Fig. 3b). Green bars: distance to the absolute fitness maximum (green dot in Fig. 3c). **b**) Same as a, but for the mutants of Fig. 3c.

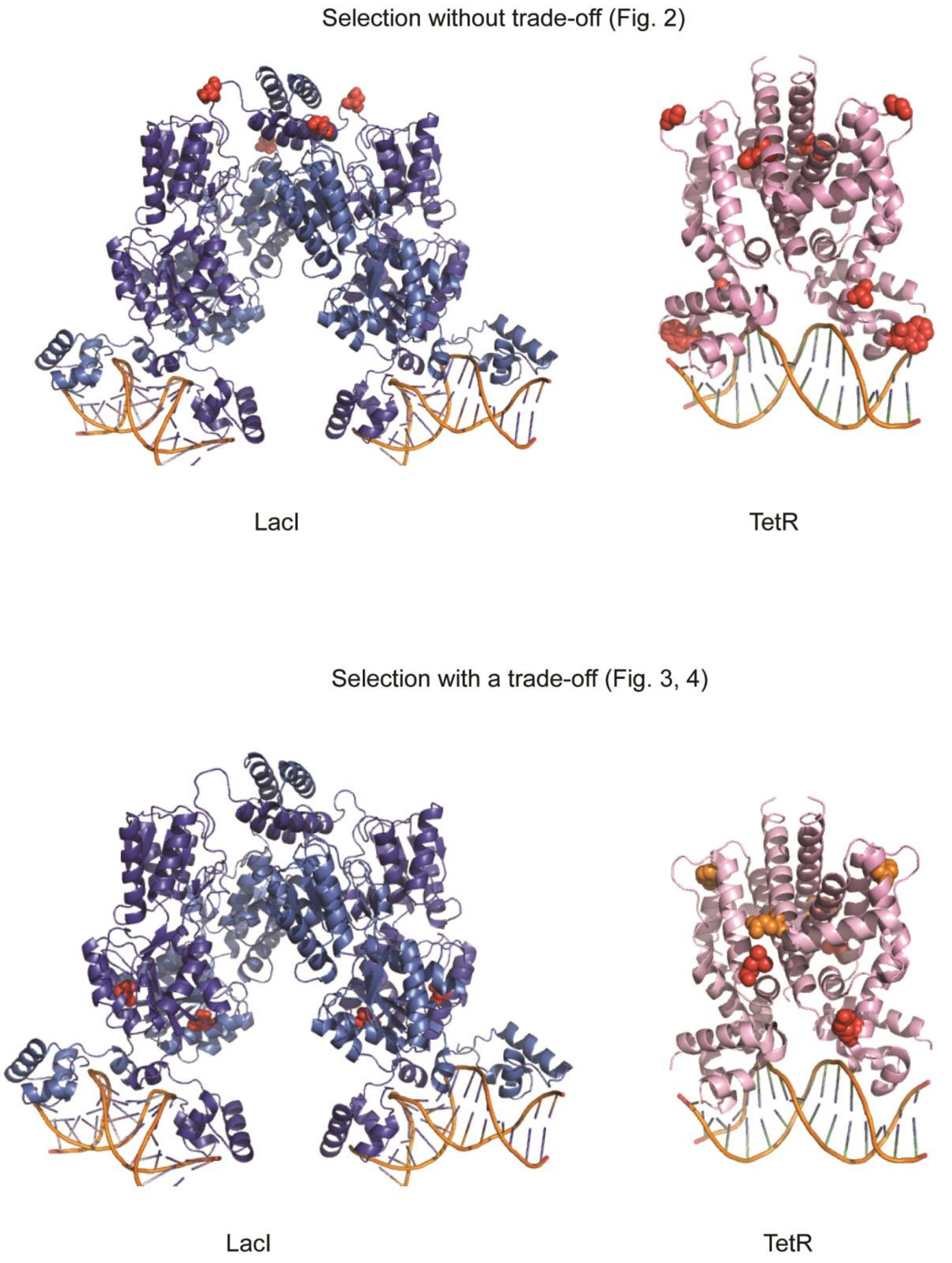


**Supplementary Fig 4**: **Mapping of mutations of onto the structures of LacI (PDB 1Z04) and TetR (PDB 1QPI) bound to their respective double-stranded operator sequences.** The mutations mentioned in supplementary Table 1 that have a consequence on the amino-acid residues are highlighted in space-filling model. Top: blue response of the mutant of Fig. 2d, also referred to as 'network M1'. Bottom: mutants of Figures 3. Red spheres are mutations obtained after the first round (red dot in Fig. 3c, also referred to as 'network M2') and orange after the second (orange dot in Fig. 3c, also referred to as 'network M3').

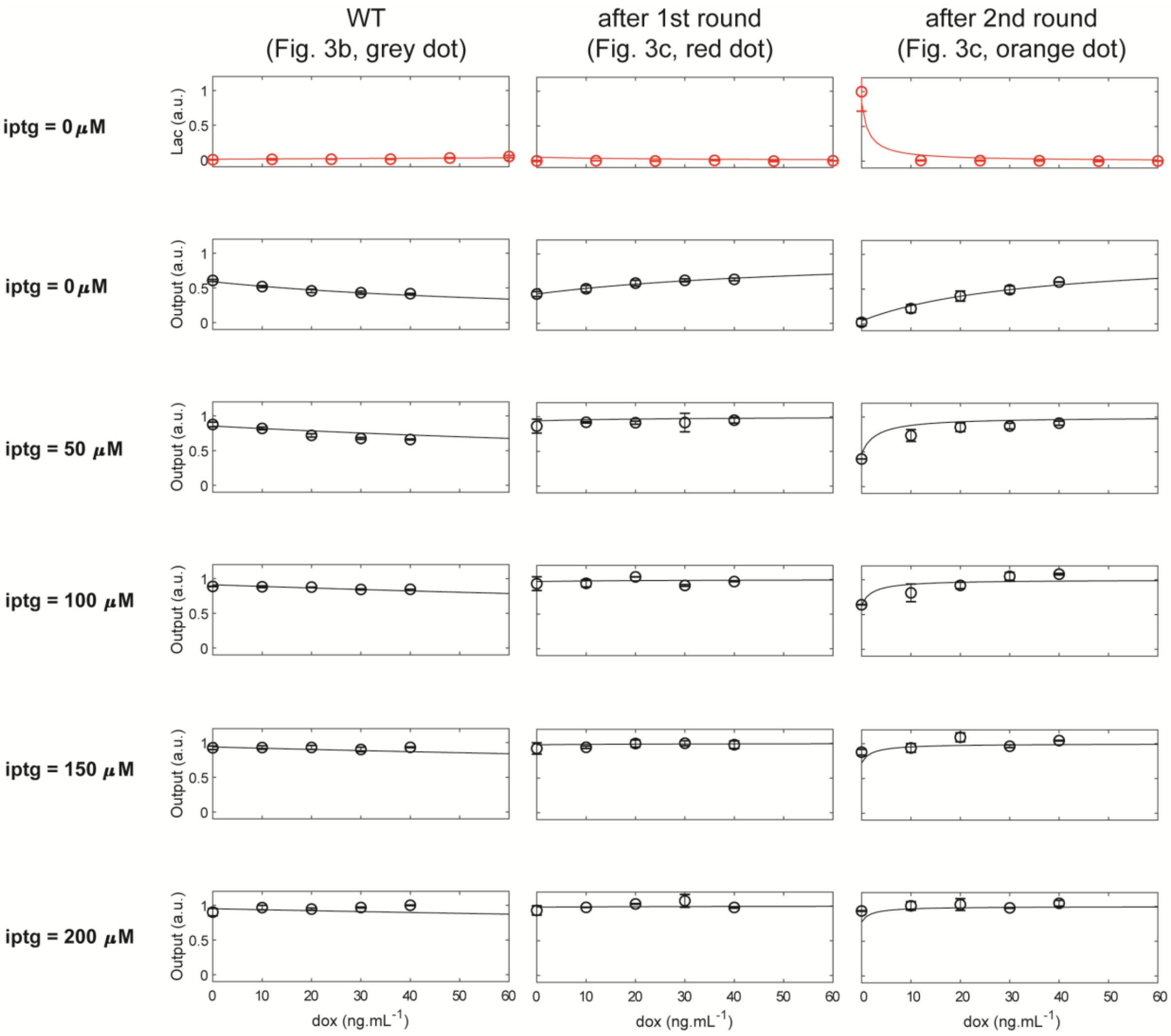


**Supplementary Fig 5. Fitting of the WT and intermediates selected after the first and second rounds of selection in Fig. 3.** Dots are experimental measurements (with standard errors over the mean) and lines the fit obtained according the Methods with the parameters of Table 1. The top row (red) are measurements of the mCherry reporter (normalized by the maximum over all experiments), which are a proxy for LacI expression levels, as a function of dox. The other rows (black) are the output level of the final cascade component (as measured with a YFP reporter).
